# Low-frequency cortical oscillations are modulated by temporal prediction and temporal error coding

**DOI:** 10.1101/076042

**Authors:** Louise Catheryne Barne, Peter Maurice Erna Claessens, Marcelo Bussotti Reyes, Marcelo Salvador Caetano, André Mascioli Cravo

## Abstract

Monitoring and updating temporal predictions are critical abilities for adaptive behavior. Here, we investigated whether neural oscillations are related to violation and updating of temporal predictions. Human participants performed an experiment in which they had to generate a target at an expected time point, by pressing a button while taking into account a variable delay between the act and the stimulus occurrence. Our behavioral results showed that participants quickly adapted their temporal predictions in face of an error. Concurrent electrophysiological (EEG) data showed that temporal errors elicited markers that are classically related to error coding. Furthermore, intertrial phase coherence of frontal theta oscillations was modulated by error magnitude, possibly indexing the degree of surprise. Finally, we found that delta phase at stimulus onset was correlated with future behavioral adjustments. Together, our findings suggest that low frequency oscillations play a key role in monitoring and in updating temporal predictions.

## 1. Introduction

Several environmental events occur regularly in time. We can take advantage of these regularities to generate temporal predictions that can enhance performance (Nobre et al., 2007; Rohenkohl et al., 2012; Vangkilde et al., 2012). For a prediction system to be successful, it is important to keep it constantly updated by monitoring when errors take place and applying the appropriate corrections. However, most temporal prediction studies have focused on situations in which there is an established temporal relation between events and little need for error monitoring and prediction updates.

Although rare in the temporal domain, several studies have investigated how our brain codes other types of prediction errors. In reinforcement learning, negative feedback has been linked to an electroencephalographic component called feedback related negativity (FRN) (Walsh and Anderson, 2012). The FRN is a frontal-central negative deflection in the event-related potential (ERP) that peaks at around 300 ms following a feedback that indicates losses or an error (Walsh and Anderson, 2012). More recently, it has been hypothesized that the FRN could be generated by perturbations in local theta band oscillations (Cavanagh et al., 2010; Cohen et al., 2007). Such perturbations are described as an increase in power and phase coherence in this frequency band in frontocentral regions (Cavanagh et al., 2010). In this view, theta oscillations would serve as a communication mechanism between brain networks, by which errors would alter oscillatory patterns and optimize the communication and the computation of relevant information (Cavanagh and Frank, 2014; Cavanagh et al., 2009). However, whether such mechanism can also be used for temporal error coding is still unknown.

As previously mentioned, the majority of studies that investigate how temporal predictions modulate performance have participants performing a task after the temporal relation between events has been learned. These studies have shown that low-frequency oscillatory brain activity (as delta, from 1 to 4 Hz) can optimize cortical excitability and enhance the processing of stimuli occurring at predicted moments (Cravo et al., 2013, 2011; Lakatos et al., 2008; Schroeder and Lakatos, 2009), as well as impair processing of temporally unexpected stimuli (Stefanics et al., 2010; van den Brink et al., 2014). Importantly, a recent study has shown that similar mechanisms seem to be involved in tasks that require a temporal judgment about the interval itself, and not just the use of the temporal information to form expectations (Arnal et al., 2014). This result supports the hypothesis that neural oscillations might serve as a possible neural mechanism for temporal predictions (Arnal and Giraud, 2012; Morillon and Barbot, 2013).

Therefore, although oscillatory mechanisms have been proposed to be important in error coding and temporal predictions, it remains largely unknown whether or not they are used when we need to learn and monitor a temporal prediction. Here, we investigated the neural mechanisms underlying violation and updating of temporal predictions. We developed a behavioral task in which participants had to monitor whether a temporal error had been made. We analyzed ERPs and oscillatory changes evoked by temporal errors in EEG recordings and investigated whether they were linked to theta oscillations. Finally, we looked for correlations between behavioral adjustment and the phase of delta oscillations.

## 2. Materials and methods

### 2.1. Participants

Twenty volunteers (18-30 years old; 11 female) gave informed consent to participate in the study. All participants had normal or corrected-to-normal vision and were free from psychological or neurological diseases. The experimental protocol was approved by the University Research Ethics Committee. Three participants did not reach the minimal performance criterion and had their data excluded from the analyses (see below for criterion of exclusion).

### 2.2. Stimuli and procedures

Stimuli were presented using the Psychtoolbox v.3.0 package (Brainard, 1997) for MATLAB on a 17-inch CRT monitor with a vertical refresh rate of 60 Hz, placed 50 cm in front of the participant. Each trial started with a fixation point. After an interval of 1.5 s, two identical audiovisual cues were presented sequentially separated by an interval of 1 second. These cues consisted of a bulls-eye (3 degrees of visual angle) presented in the center of the screen and an auditory tone (1000 Hz, 70 dB) both presented for 100 ms.

A third stimulus (which we refer to as the target) was an auditory tone (500 Hz, 70 dB) presented for 100 ms. The temporal onset of this target was controlled by the participants. Their main task was to generate the target (tone) at an expected time point by pressing a button while taking into account the inserted delay between their press and the target occurrence. Participants were instructed that the interval between the second cue and the target should be identical to the interval between the first and second cues. Therefore, in order to produce the target at the correct moment, they had to consider the delay between their action and target presentation. Participants were told that the interval between the two cues was constant throughout the experiment, but the exact interval value (1 s) was never mentioned (Figure 1A).

**Figure 1:**
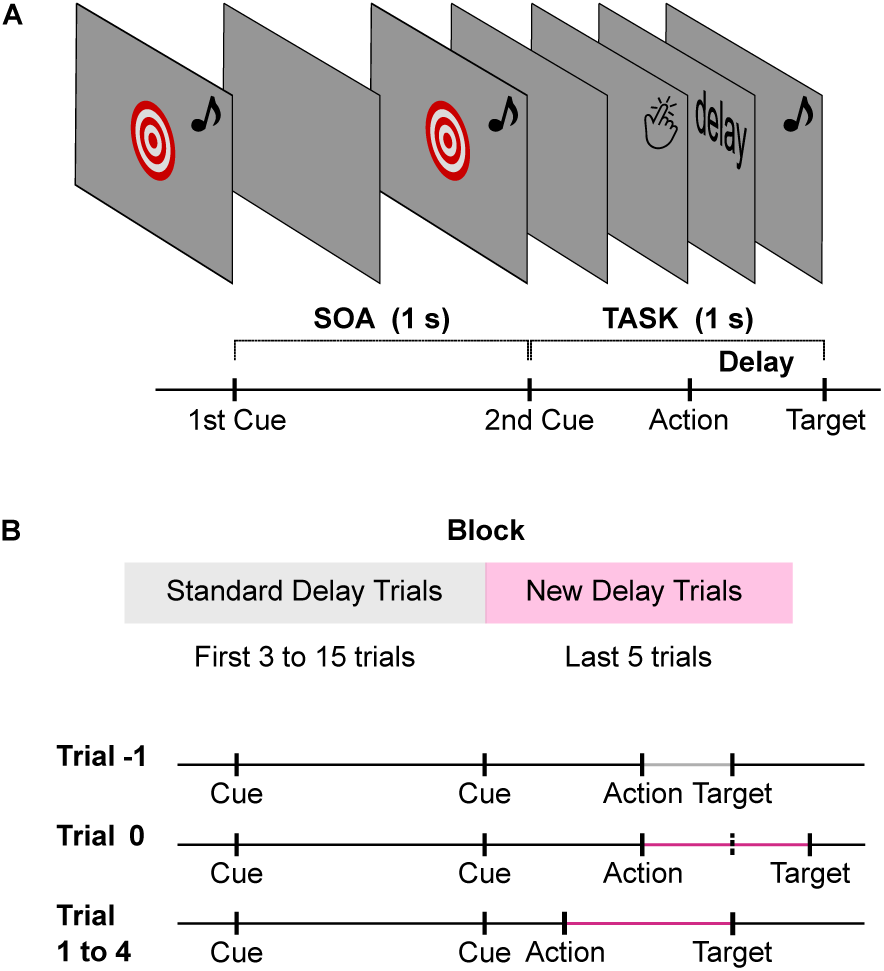
Schematic illustration of task structure. A) The main task was to generate a third stimulus (target) at an expected time point, by pressing a button while taking into account the delay between the act and target occurrence. Participants were instructed that the interval between the second cue and the target should be identical as the interval between the first and second cues. B) All blocks started with trials where the action-target delay was 50 ms (standard delay trials). At a given trial, a new delay (between 300 ms and 700 ms) was inserted and kept constant for five trials. In trial −1 (before the new delay is inserted), participants are adapted to the delay of 50 ms between their action and outcome and perform the button press at the appropriate moment. In trial 0 (when the new delay is inserted), the outcome comes later than expected. Based on this error, in the following trials (1 to 4) participants have to update their temporal prediction and anticipate their action.

A delay was inserted between each button press and the occurrence of the target. In the first few trials of each block, the delay between action and target was 50 ms (standard delay). However a new delay between 300 ms and 700 ms was inserted in a given trial within the block and remained fixed for five trials. The change in the action-target delay was intended to cause a temporal prediction violation and have the target appear latter than expected by the participant. Once the new delay had been inserted, participants had to update their temporal model and anticipate their action in order to make the target appear at the appropriate moment in the remaining trials of the block. After the five trials with the new delay had been presented, that particular block ended. Thus, each experimental block started with standard delay trials and ended with five new delay trials (Figure 1B).

Participants were informed that the action-target delay would change in a given trial and remain fixed for five trials, after which the current block would end. Importantly, they could not predict when or by how much the delay would change, as the new delay could be inserted randomly between the 4th and 15th trial in each block. Moreover, a change in delay was made only if participants absolute errors in the previous three standard delay trials were smaller than 100 ms (i.e., if the target appeared between 900-1100 ms after the second cue). If the participant did not reach this criterion until the 15th standard delay trial, a new delay was inserted in the 16th trial. These two rules inhibited behavioral anticipation to the new delay.

Participants who failed to perform well with the standard delay for more than 10% of the blocks were excluded from the analyses. Blocks in which the participant did not reach the performance criterion until the 15th trial were excluded from both behavioral and EEG analyses. Temporal errors over 1.5 seconds were considered omission errors and removed from subsequent analyses (three trials in total, one omission error for three different participants). Importantly, explicit feedback about the participants performance was given only at the end of each experimental block. Therefore, no information about errors was shown throughout a block and participants could only extract information about their performance based on their own temporal predictions. Each session consisted of 50 blocks, and lasted between 40 to 60 min. Each block consisted of 8 to 20 trials. Participants underwent two blocks of practice trials before the experimental session began.

### 2.3. EEG recordings and pre-processing

EEG was recorded continuously from 64 ActiCap Electrodes (Brain Products) at 1000 Hz by a QuickAmp amplifier (Brain Products). All sites were referenced to FCz and grounded to AFz. The electrodes were positioned according to the International 10-10 system. Additional bipolar electrodes registered the electrooculogram (EOG).

EEG pre-processing was carried out using BrainVision Analyzer (Brain Products). All data were down-sampled to 250 Hz, re-referenced to the averaged earlobes, and epoched from −3500 ms to 2000 ms relative to targets onset. An independent component analysis (ICA) was performed on filtered data (0.5 Hz to 30 Hz) to reject eye movement artifacts and the resulting component coefficients were applied to the unfiltered data. Eye-related components were identified by comparing individual ICA components with EOG channels and by visual inspection. The number of trials rejected for each subject was small (an average of 1.3% of trials across subjects, maximum of 4.7%). For the feedback related negativity (FRN) analyses, continuous EEG was filtered off-line with a 1-30Hz band-pass filter (24 db/oct) and epoched from −250 and 700 ms relative to target presentation.

Time-frequency analyses were performed on unfiltered data using the SPM8 and Fieldtrip toolbox for MATLAB (Oostenveld et al., 2011). Individual frequency bands (delta, 0.5-4 Hz; theta, 4-8 Hz; alpha, 8-16 Hz; low-beta, 16-24 Hz; high-beta, 24-32 Hz) were extracted using third-order dual-pass Butterworth filters, and the phase and power of these narrow-band signals were calculated using the Hilbert transform (Besle et al., 2011; Kayser et al., 2009; Ng et al., 2012a,b). For delta phase, the Butterworth filters were applied to the continuous data, while for other frequencies the filter was applied to the epoched data (-3500 ms to 2000 ms relative to target presentation). Power was defined as the squared absolute value, and phase was defined as the phase angle of the Hilbert signal. To characterize the phase distribution across trials, we calculated the intertrial coherence (ITC) for each frequency and time point (Lachaux et al., 1999; Tallon-Baudry et al., 1996).

For each frequency band, electrodes of interest were chosen based on the grand-averaged activity for trials 0 to 4. Cluster-based analyses were performed by calculating permutation tests in which experimental conditions were intermixed within each subject in 1000 random permutations (Oostenveld et al., 2011).

### 2.4. Statistical analyses

In the majority of analyses, the dependent variable of interest was submitted to a repeated-measures ANOVA with trial as the main factor. When there was a significant effect of trial, a multiple contrasts procedure, as proposed by Scheffé’s, was used to compare selected contrasts (Zar, 1999). Specifically, the following contrasts were tested: (1) if the measured value in trial 0 was different from the average of trials 1 to 4; (2) if the measured value in trial 1 was different from the average of trials 2 to 4; (3) if the measured value in trial 2 was different from the average of trials 3 and 4; (4) if the measured value in trial 3 was different from trial 4. The tested contrasts were selected to measure if and when the dependent variable of interest stopped changing as a function of trial. For example, if learning is instantaneous, one would expect a significant main effect of trial and only the first contrast to be significantly different from zero. If, on the other hand, learning keeps taking place until the end of the block, one would expect a significant main effect of trial and all contrasts to be significant.

## 3. Results

Results are presented and discussed separately for each of the three main goals: 1) to describe behavioral adjustments following a temporal error; 2) to identify whether temporal errors evoke activity related to error processing (as the FRN and theta oscillations); and 3) to determine whether behavioral adjustments are biased by delta oscillations.

### 3.1. Behavioral Results

The main dependent variable for the behavioral analysis was the error committed by participants following a change in the action-target delay. This error was calculated with signed and absolute values in separate analyses. In both cases, we calculated the difference between the moment the target should have been presented (1s after the second cue) and the moment it actually was presented, where zero represents perfect performance. For signed errors negative/positive values indicate that the target was presented earlier/later, corresponding to anticipated or delayed action, respectively. We used both types of analyses to investigate whether results were due to positive and negative errors canceling each other out. All of our analyses focuses on the five trials with the new inserted delays in each block (which we refer to as trial 0 to trial 4). Trial 0 represents the first trial with new delay and trial 4 the last trial with that delay.

For the signed errors, average error for trials 0 to 4 were submitted to a repeated-measures ANOVA with trial as a factor. There was a significant effect of trial (F(4,64) = 302.38, p <0.001; Figure 2A). We applied Scheffé’s method on the four multiple contrasts previously described. Error in trial 0 was significantly larger than in subsequent trials (p <0.001), while no other contrasts were statistically significant (p >0.9). We performed a similar analyses to the averaged absolute errors and found a similar pattern: there was a main effect of trial (F(4,64) = 350.84, p <0.001), with absolute errors in trial 0 being significantly larger than errors in subsequent trials (p <0.001). Likewise, no other contrasts were statistically significant (p >0.4). These results suggest that participants quickly adjusted performance after a single exposure to the new delay.

**Figure 2:**
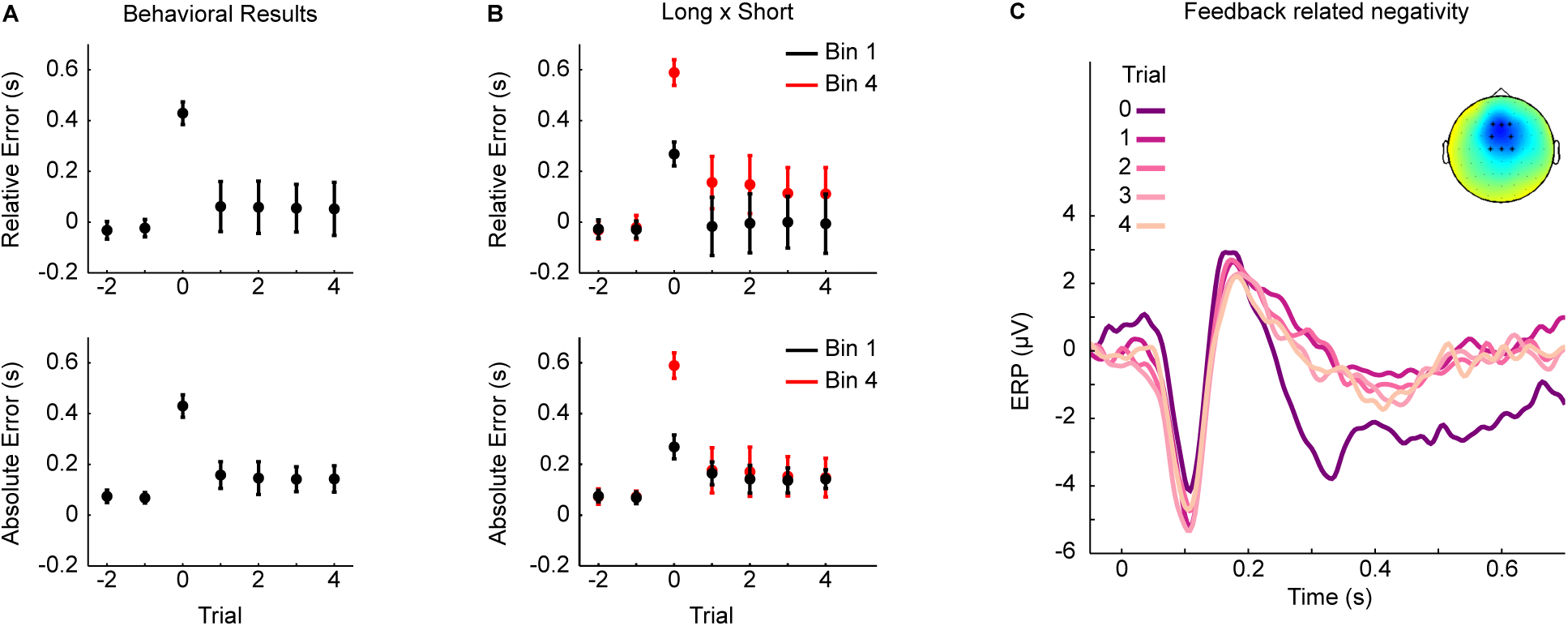
A) Temporal error (mean ± standard deviation) as a function of trial. The two last standard delay trials of each block are represented by negative values in the x-axis. The first trial with the new delay is represented as trial 0, while the remaining trials with the new delay are represented by positive values in the x-axis. Upper panel shows the relative error and the lower panel the absolute error. B) Temporal error (mean ± standard deviation) for the shortest and longest delay bins as a function of trial. C) Event-related potential evoked by the target (zero on the x-axis indicates target onset). The number of the trial with the new delay is color coded, with darker/lighter colors representing the first/last trials with the new delay. The inset shows the topography of the Feedback Related Negativity, for the period between 200 ms and 400 ms after target onset.

To confirm that participants were indeed using information from the previous trial to update their actions, we analyzed how the delay on trial 0 in each block modulated the time of their action on the subsequent trial. A linear regression between delay on trial 0 and time of action on trial 1 was estimated for each participant, and slope estimates were submitted to a one-sample t-test. We found that the slopes were significantly smaller than zero (average slope = −0.43, t(16) = −5.99, p <0.001), indicating that larger delays in trial 0 were followed by larger anticipations of action.

Next, we investigated the effect of delay magnitude on temporal update. Given that the inserted delay could vary from 300 ms to 700 ms, we investigated if the magnitude of the delay influenced learning rate and final performance in each block. We binned delays in four quantiles, each with 25% of the data (midpoints 0.35 s, 0.46 s, 0.59 s, and 0.67 s,) and performed a similar analysis as described above (Figure 2B). For the signed errors, all delay bins had a similar learning pattern with a main effect of trial (Bin 1: F(4,64) = 66.61, p <0.001; Bin 2: F(4,64) = 160.31, p <0.001; Bin 3: F(4,64) = 232.76, p <0.001; Bin 4: F(4,64) = 300.07, p <0.001). Scheffé’s method on the four contrasts indicated that in all bins the average signed error in trial 0 was larger than in the subsequent trials (p <0.001), while no other contrasts were statistically significant (p >0.2). For the absolute errors, we found the same learning pattern. There was an effect of trial for all delay bins (Bin 1: F(4,64) = 34.35, p <0.001; Bin 2: F(4,64) = 191.92, p <0.001; Bin 3: F(4,64) = 296.86, p <0.001; Bin 4: F(4,64) = 355.31, p <0.001).. For all bins, error in trial 0 was larger than the error of the subsequent 4 trials (p <0.001), while no other contrasts were statistically significant (p >0.2).

To compare final performance for the different delays, the average error at trial 4 for each bin was submitted to a repeated-measures ANOVA with Delay as a factor. For signed errors, there was an effect of delay on final performance (F(3,48) = 20.04, p <0.001). Pairwise post-hoc analyses (Tuckey correction) indicated that average error for the shortest delays (Bin 1 and Bin 2) were significantly different from average errors of longest delays (Bin 3 and Bin 4), p <0.05. Average errors between Bin 1 and 2; and between Bin 3 and 4 were not significantly different (p >0.1). For absolute errors, however, there was no significant effect of delay (F(3,48) = 0.27, p = 0.85). These results suggest that the magnitude of the inserted delay modulated participants’ final performance.

### 3.2. Target-related activity

In a first step, we investigated broadband ERPs (1 Hz to 30 Hz) elicited by the target. Data were epoched from −250 and 700 ms relative to target presentation and averaged separately for trials 0 to 4.

There was a stronger negative component in trial 0 evoked in frontocentral electrodes (F1/Fz/F2, FC2/FC1, C1/Cz/C2) compared to other trials. To analyse whether temporal errors elicited a FRN, we calculated the average activity in these electrodes for the time period between 200 ms and 400 ms after target’s presentation for each trial. A repeated-measures one-way ANOVA indicated a significant modulation of activity as a function of trial (F(4,64) = 9.56, p <0.001). A similar multiple contrast showed that activity in trial 0 was different from subsequent trials (p <0.001) with no other contrast reaching significance (p >0.77). To show the topography of this effect, we subtracted the mean activity of trial 0 from the averaged activity of trials 1 to 4 (Figure 2E). The topography and time period of the effect are similar to previous reports of FRN (Miltner et al., 1997; Walsh and Anderson, 2012).

Given the strong relation between FRN and theta oscillations, we looked into theta activity produced by the target. Targets evoked a strong theta power in frontocentral electrodes, as commonly found in studies that investigate theta oscillations and error coding. However, a one-way ANOVA with cluster correction found no periods with a significant modulation of theta power as a function of trial (cluster stats 2.64, cluster p >0.9).

We found that targets evoked a high theta phase reset, as measured by ITC, in similar frontocentral electrodes (Figure 3A). A cluster analysis indicated a significant modulation of theta ITC by trial in the period between 220 ms to 380 ms (cluster stats 25.87, cluster p <0.05). Scheffe´’s method on the multiple contrasts at the cluster period indicated that theta ITC in trial 0 was significantly larger than other trial’s ITC (p <0.05). No other contrasts were statistically significant (p >0.8).

**Figure 3:**
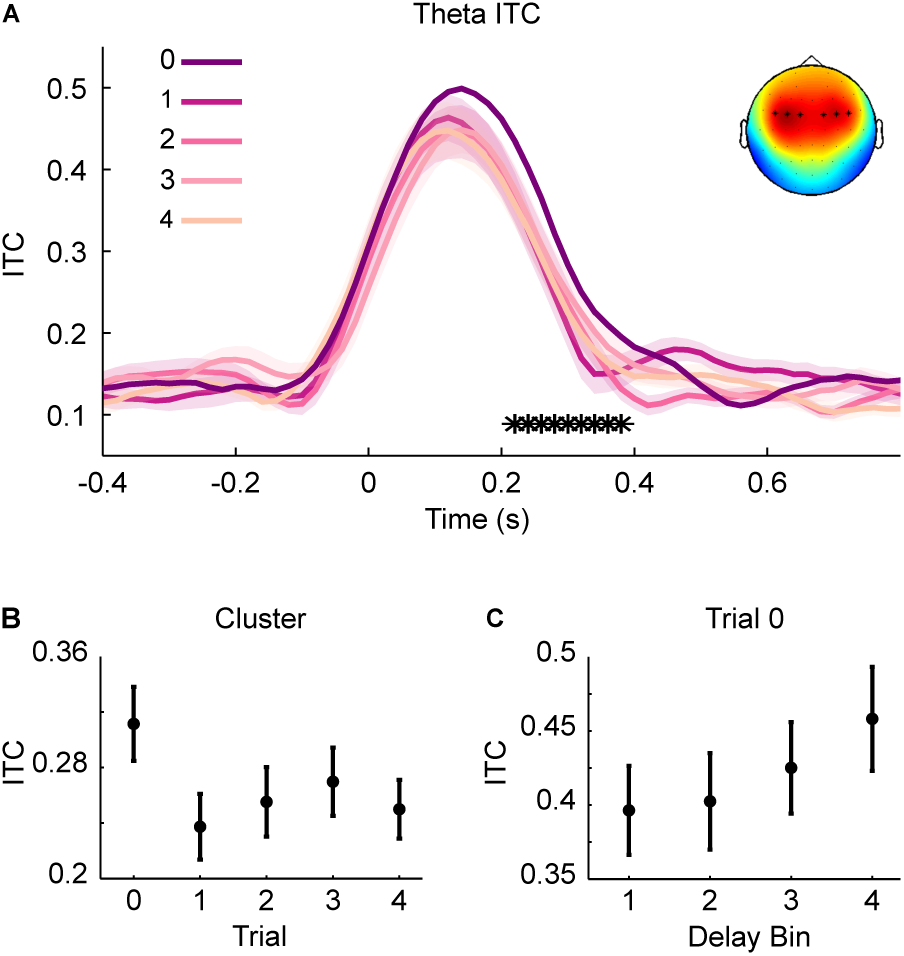
A) Theta ITC evoked by target (zero on the x-axis indicates target onset). Asterisks indicate the significant temporal cluster. The topography shows the electrodes with strong theta ITC. The number of the trial with the new delay is color coded, with darker/lighter colors representing the first/last trials with the new delay. B) Average theta ITC during the cluster period (220 ms to 380 ms) as a function of trial (mean ± S.E.M). C) Theta ITC (mean ± S.E.M) as a function of binned delays in trial 0 during the cluster period. Bin 1 corresponds to shortest delays whereas Bin 4 corresponds to longest action-target delays.

To investigate whether theta ITC was higher as a function of the magnitude of the new delay, we performed a follow-up analysis focusing on the cluster period but only on the first trial with the new delay of each block (trial 0). Notice that, in these trials, participants could not foresee that a larger delay would take place, so they did not anticipate their actions accordingly. Because ITC cannot be measured in single trials, we performed a similar binning procedure as for the behavioral analyses and for each bin calculated theta ITC. Next, we calculated a Spearman rank order correlation between delay bin and theta ITC for each participant. At the group level, participants Spearman correlations were compared to zero using a one-sample t-test. We found that longer delays were significantly associated with stronger theta ITC (Rhos = 0.38 0.13, t(16) = 2.99, p <0.01).

### 3.3. Anticipatory activity: delta oscillations

Given the known effects of delta phase being modulated by temporal expectations, we investigated whether delta phase was involved in temporal prediction and adjustment. There was a high ITC of delta phase at target presentation in left-central electrodes (C5/C3, CP3/CP5, Figure 4A), similar to previous results of delta phase effects on temporal processing (Arnal et al., 2014; Stefanics et al., 2010). We investigated whether delta phase was concentrated around different angles as participants adapted to the delay between their action and outcome. Delta phase for the period between −50 ms and 0 ms around target onset was averaged for each participant. Because phase is a circular measure, to use regular statistical analyses we first performed a normalization step for each participant, by subtracting the average mean of the trial 4 from all other trials (using circular distance). With this subtraction, it was possible to test if targets became more concentrated around a specific phase as participants adapted to the new delay. Our dependent variable was the average distance (in degrees) between delta phase in each trial from the phase where the target was presented when participants were adapted to the delay. These average distances were submitted to a repeated measures one-way ANOVA and there was a significant modulation by trial (F(4,64) = 22.531, p <0.001). Average distance in trial 0 was significantly larger than other trials (p <0.001), while no other differences were statistically significant (p >0.5), suggesting that, as participants learned the new delay between their action and target, brain states were realigned, resulting in the target being consistently presented in a specific delta phase (Figure 4B and 4C).

**Figure 4:**
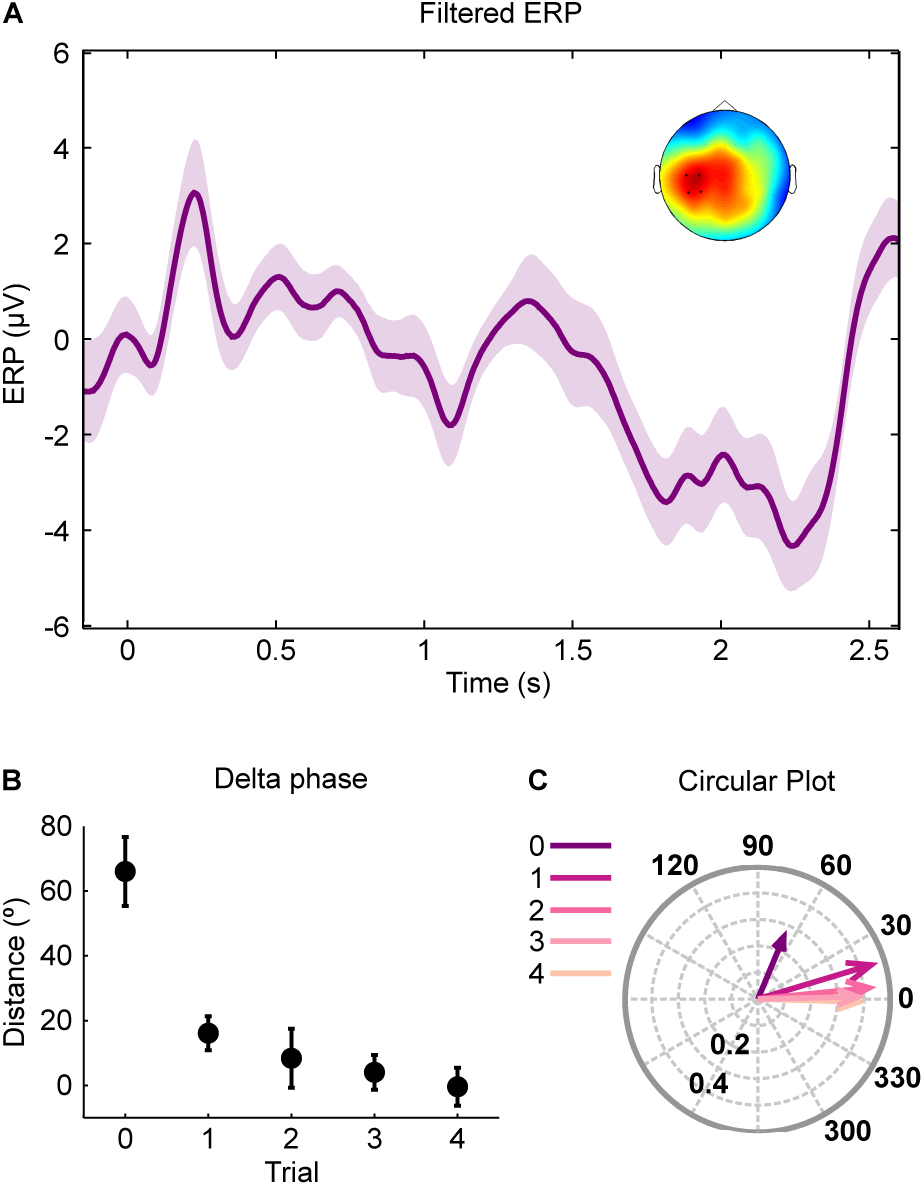
Delta Phase. A) ERP waveforms filtered between 0.5-4 Hz. The zero label on the time axis represents first cue presentation, 1s represents second cue presentation and 2s represents the moment target should have been presented. Here, we plotted ERP from trials 0 in which targets appeared after at least 2.55 s. The topography (inset) illustrates the electrodes that showed higher delta ITC at target presentation (C5/C3, CP3/CP5). B) Distance of delta phase to average phase at trial 4. C) Circular plot illustrating both distance of delta phase to average phase at trial 4 and ITC. Distance is illustrated by the angle and delta ITC is represented by arrows magnitude. Circles inside the plot indicate the ITC scale. The number of the trial with the new delay is color coded, with darker/lighter colors representing the first/last trials with the new delay.

To investigate further the relation between delta oscillations and temporal prediction, we analyzed whether delta phase on target onset in trial 0 helped to predict the adjustment participants made in the subsequent trial. If delta phase reflects temporal expectations about when the target will be presented, then having access to this information should improve our prediction of the adjustments participants will make on the following trial.

We fitted two mixed linear models to determine whether the moment of action in trial 1 could be explained better by taking into account delta phase on tar-get onset on trial 0. Specifically, we compared the fol-lowing models:

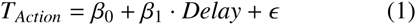

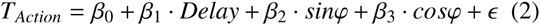

where *T*_*Action*_ is the moment of the participants action on trial 1, Delay is the physical interval between action and target presentation, and β_*i*_ represents the fitted coefficients. In the first model, only the delay between action and target was used to fit the moment of action of participants in the following trial. In the second model, delta phase at target onset in trial 0 was added.

The fits of the two models were compared using Akaikes information criterion (AIC) and a likelihood ratio test. We found that the inclusion of delta phase significantly improved the fit (no-phase model AIC = −668.19, phase model AIC = −672.19; log-likelihood ratio test = 12.33, df = 2, p <0.01). This suggests that not only the delay, but also delta phase at target onset in trial 0 significantly modulated participants action in the following trial (Figure 5).

**Figure 5:**
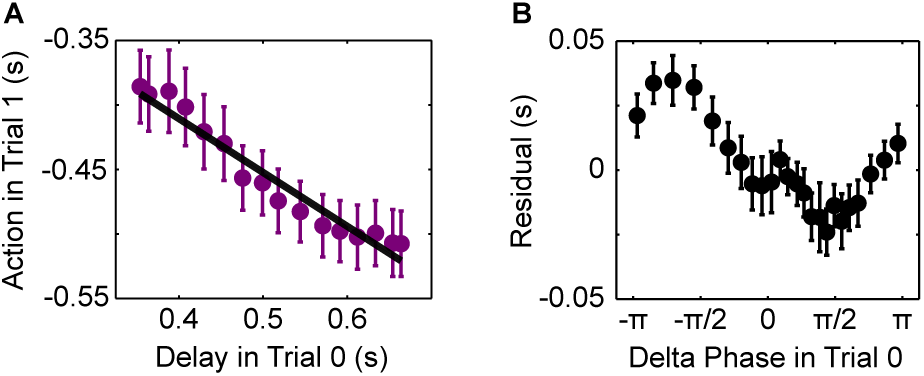
Model comparison. A) Behavioral adjustment in trial 1 as a function of delay in trial 0, fitted by the mixed linear model without delta phase (Eq. 1). B) Residuals of the first model plotted as a function of delta phase at target presentation in trial 0.

We also looked at delta power from the same electrodes. At target presentation, there was a significant modulation of delta power as a function of trial (F(4,64) = 3.73, p <0.05). Scheffé’s method on the four contrasts indicated that delta power on trial 0 was larger than in subsequent trials (p = 0.015). No other contrasts were statistically significant (p >0.8). However, delta power in the first trial was not modulated by delay magnitude (mean Rho = −0.074, t(16) = −1.71, p >0.05).

### 3.4. Statistical analyses of other frequency bands

We performed similar cluster-based analyses on alpha and beta power. A decrease in alpha power was observed after target presentation in parietal sites (Pz/P2/P4/PO4/POz). A cluster analysis showed that this decrease was modulated by trial in the period from −180 ms to 540 ms around target presentation (cluster stats 185.71, cluster p <0.001). Scheffé’s method on the four multiple contrasts indicated that alpha power in trial 0 was significantly smaller than the average activity from all other trials (p <0.001), while no other differences were significant (p >0.5). We investigated further to test whether alpha power in the first trial was modulated by delay, in an analysis similar to theta ITC. There was no significant correlation between delay and alpha power (mean rho = −0.20, t(16) = −1.78, p >0.05).

For both low and high beta power, there was a small increase prior to target presentation at the grand averaged activity in fronto-central electrodes (F1/Fz,FC2/FCz/FC1/FC3, C3/C1). However, no significant modulation of beta power by trial was found.

## 4. Discussion

In the present study, we investigated the mechanisms underlying violation and updating of temporal predictions. We found that participants are able to adapt quickly to a new temporal relation between their action and outcome. Temporal errors elicited electrophysiological markers classically related to error coding. These markers were modulated by the magnitude of the error. Finally, we showed that delta phase at the moment of target onset is correlated to future behavioral adjustments.

Several studies have investigated how humans and non-human animals learn and adapt to new temporal relations between events (Church, 2014; Reyes and Buhusi, 2014). As in previous studies, there was a rapid temporal learning of a new delay (Balsam and Gallistel, 2009; Higa, 1997; Simen et al., 2011). A single exposure was sufficient for participants to adjust their future actions and such a rapid and accurate learning suggests a precise error controlling mechanism.

The electrophysiological results revealed that temporal errors elicited signals normally related to error processing. It has been previously suggested that these signals, frontal theta oscillations and the feedback related negativity, are related and may serve as possible mechanisms for cognitive control. They have been found in situations of error, conflict, punishment and novelty (Cavanagh and Frank, 2014). Previous studies focusing on the relation between error signals and temporal processing have used tasks with explicit feedback (Miltner et al., 1997; van de Vijver et al., 2011). In the current study, we have shown that an incorrect temporal prediction by itself, without an explicit feedback, can lead to an increase in these error-related components, especially for intertrial phase coherence in the theta band. Frontal medial theta phase reset has been suggested to be a biologically plausible candidate for neuronal computation and communication (Cavanagh and Frank, 2014; Cavanagh et al., 2011). According to this hypothesis, the need for cognitive control can instantiate transient functional networks across spatially distal sites by means of phase alignment. In our results, larger temporal errors elicited stronger phase reset, possibly coding the degree of surprise caused by the new delay (Cavanagh et al., 2011; Hauser et al., 2014; Talmi et al., 2013).

Together, our behavioral and electrophysiological results suggest a precise error monitoring mechanism for temporal processing. A remaining question is how temporal predictions are represented cortically. Recent proposals have suggested that delta oscillations might play an important role in temporal predictions (Arnal et al., 2014; Arnal and Giraud, 2012). In a recent study, Arnal and colleagues found that delta phase correlated with response accuracy in a temporal judgment task (Arnal et al., 2014). However, in their study participants made a simple judgment about whether or not a tone was delayed with regard to an established rhythm. In our study, temporal information could not be used to form a simple categorical response. On the contrary, participants had to track time continuously to perform the task. Moreover, the moment of action in the following trial is a more sensitive measure of participants temporal perception. Our results showed that delta phase was correlated with participants action in the subsequent trial. This suggests that delta oscillations might reflect the temporal prediction of when the target will be presented. It is not possible, however, to determine whether delta phase is driving the prediction itself or if it is a downstream consequence of higher areas tuning to the temporal regularities of the task.

In both our and Arnals study, the task had an underlying rhythm in the delta band. In fact, most studies that have shown effects of delta phase on performance had an underlying temporal structure in this frequency band (Besle et al., 2011; Cravo et al., 2013; Schroeder and Lakatos, 2009). However, some studies have also shown a correlation between delta oscillations and temporal processing in tasks without an external rhythm (Laubach et al., 2015; Stefanics et al., 2010). In both cases (presence or absence of a rhythm), delta oscillations seem to be realigned such that relevant information falls within a similar phase. When a temporal prediction is wrong, targets will fall in different delta phases, which can be used, in turn, as a measure of error, resulting in precise adjustments (Arnal et al., 2014).

Although our results are in agreement with the general finding that delta oscillations are important in temporal expectations, other frequency bands have also been hypothesized to work as predictive timing mechanisms. For example, it has been found that beta oscillations are related to temporal expectations both by its increased amplitude (Arnal, 2012; Fujioka et al., 2012; Kilavik et al., 2013) and by its coupling with delta phase (Arnal et al., 2014). However, we did not find any significant evidence of beta oscillations being modulated by temporal expectation or temporal error. Contrary to delta oscillations, which have been found both in tasks with and without an underlying rhythm, the relation between beta oscillations and temporal expectation has been found more consistently in tasks with rhythmic sequences. Thus, it is possible that our experiment was not ideal for this kind of modulation. Moreover, it is still unclear how different rhythms modulate beta power (Meijer et al., 2016).

Although no other differences were seen in higher frequency bands, we did observe a significant alpha power suppression in trial 0. However, contrary to previous findings, we did not find that alpha power suppression was correlated with delay magnitude (Arnal et al., 2014). One possibility for this difference is that our task did not require an explicit judgement about the temporal interval, suggesting that the relation between alpha suppression and time can be task dependent.

In conclusion, we have found that cortical oscillations are correlated with monitoring and updating of temporal predictions. Temporal errors elicited known oscillatory correlates of error. These findings suggest that rhythmic changes in cortical excitability might reflect temporal predictions which in turn can be used for behavioral adjustments.

## 5. Acknowledgments

This work was supported by São Paulo Research Foundation (FAPESP), grants 2013/24889-7 and 2014/08389-7. The authors would like to thank the members of the Timing and Cognition Laboratory at UFABC (http://neuro.ufabc.edu.br/timing/) for useful discussions and suggestions on earlier manuscript versions; and Felipe Cardoso for helping with pilot experiments.

